# Pericyte-Derived Cancer-Associated Fibroblasts Correlate with Poor Survival and Are Enriched After Chemoradiotherapy in Glioblastoma

**DOI:** 10.64898/2026.07.02.736148

**Authors:** Aly Ismailov, Maria Poptsova

## Abstract

The role of cancer-associated fibroblasts (CAFs) in glioblastoma remains unclear, as their existence in the brain tumor microenvironment is still debated, given that the normal brain parenchyma is devoid of fibroblasts. It is unclear whether cells described as CAFs represent a distinct stromal population or a transcriptional state of perivascular cells such as pericytes. The aim of this study was to determine the identity, origin, and functional relevance of CAFs in glioblastoma. We analyzed 54 single-cell RNA sequencing datasets together with 88 bulk RNA sequencing samples. We identified a continuous transcriptional spectrum linking endothelial cells, pericytes, and CAFs, supporting pericytes as the most likely source of CAFs in glioblastoma. We further derived and validated robust CAF- and pericyte-specific gene signatures, enabling clear separation of these populations across cohorts. Reproducible CAF-associated ligand–receptor interactions were enriched in angiogenesis and immune modulation pathways. In bulk RNA-seq data, both CAF signature scoring and deconvolution consistently demonstrated increased CAF abundance in IDH-wildtype gliomas and further enrichment after chemoradiotherapy, while selective CYP1B1 expression in CAFs suggested a potential association with therapy-induced tumor adaptation. Overall, CAFs represent a distinct, pericyte-related stromal population in glioblastoma with conserved transcriptional and signaling programs. High CAF signature scores were associated with poorer overall and progression-free survival and were enriched in IDH-wildtype and post-chemoradiotherapy gliomas, suggesting a role for CAFs in therapy-associated remodeling of the tumor microenvironment in aggressive disease.

## Introduction

Glioblastomas (GBM) are the most common primary brain tumors, characterized by high mortality and an abysmal prognosis. Moreover, they account for approximately 16% of primary central nervous system neoplasms. Despite aggressive treatment strategies, including surgical tumor resection followed by radiotherapy and chemotherapy, the survival rate of GBM patients has remained stagnant at 12–18 months for many years [1]. Given the limited success of current therapies, there is a pressing need for a more detailed investigation of the cytological and biological properties of these tumors.

Cancer-associated fibroblasts (CAFs) have been identified in numerous cancer types [2], where their central role in maintaining the tumor microenvironment (TME) is well established. However, because fibroblast populations are absent in the native brain, it was long assumed that CAFs are also absent in GBM. The advent of single-cell technologies has challenged this view, with recent reports describing cell populations expressing fibroblast markers within GBM [3–6]. Nevertheless, this topic remains largely unexplored, particularly regarding the progenitor cells that give rise to CAFs, their cellular origins, and their functional role in the GBM TME.

Several factors may contribute to this uncertainty, including potential inter-patient heterogeneity, the relatively low abundance of CAF-like cells within the overall tumor mass, and technical limitations of single-cell RNA sequencing, where enzymatic tissue dissociation may lead to selective loss or underrepresentation of fragile or rare stromal populations [7]. In addition, evidence from other tumor types suggests that CAFs may arise from diverse cellular sources, and candidate progenitor populations include quiescent fibroblasts, endothelial cells, pericytes, stellate cells, bone marrow-derived mesenchymal cells, adipocytes, and even glial-lineage cells [1, 8–13]. Consequently, despite growing interest in tumor stromal biology, the cellular origins of CAFs in remain unresolved.

Here, we systematically characterized cancer-associated fibroblasts in GBM using integrated analysis of single-cell and bulk RNA sequencing datasets. We focused on defining their cellular origin, transcriptional programs, intercellular communication networks, and clinical relevance across molecular subtypes and treatment states.

## Materials and Methods

### Single cell analysis

The discovery cohort was based on dataset GSE173278 [14], while the validation cohort was generated by integrating publicly available datasets GSM3828672 [15], GSE103224 [16], GSE135045 [17] and 10x Genomics GBM 3’ v3 scRNA-seq dataset. Standard single-cell RNA-seq preprocessing and quality control were performed for all datasets using Seurat R package. Cells were filtered based on mitochondrial content (<20% mitochondrial gene expression), minimum gene detection per cell (≥300 genes), and minimum gene expression per gene (≥3 cells). Potential doublets were identified using DoubletFinder and removed prior to downstream analysis. RNA contamination was assessed using DecontX, and corrected where applicable. Dimensionality reduction was performed using principal component analysis (PCA), followed by construction of a shared nearest-neighbor graph and unsupervised clustering using the Louvain algorithm. Uniform manifold approximation and projection (UMAP) was used for visualization. Batch effects across datasets were corrected using Harmony integration. Malignant cells were identified using InferCNA-based inference of large-scale copy number variation profiles [18], while non-malignant cell populations were annotated based on canonical marker gene expression. Differentiation trajectories were reconstructed using diffusion maps and diffusion pseudotime with default parameters as implemented in the R package destiny. Trajectory inference and generalized additive model (GAM) analysis were performed independently for the discovery and validation cohorts. Within each cohort, a GAM was fitted to each gene to identify genes exhibiting significant expression changes along the ingerred trajectory. Genes with Benjamini–Hochberg adjusted p-values < 0.05 were considered dynamically expressed and retained for downstream analysis and visualization.

### Identification of CAF and pericyte gene signatures

Gene signatures for CAFs and pericytes were derived using an iterative differential expression analysis strategy. Differential expression analysis was performed using the FindMarkers function from the Seurat package with the following thresholds: log fold change > 0.25, adjusted p-value < 0.05, and a minimum fraction of expressing cells > 0.1. Briefly, each target cell population (CAFs or pericytes) was compared individually against every other cell population, including malignant glioblastoma subtypes (MES1-like, MES2-like, NPC1-like, NPC2-like, OPC-like, AC-like, and glioblastoma stem cells - GSCs), to distinguish stromal markers from genes associated with the mesenchymal transcriptional program of glioblastoma, as well as against non-malignant cell types. Genes consistently identified as differentially expressed in the target population across all pairwise comparisons were retained to define a preliminary gene signature. The same procedure was independently applied to the validation cohort. To ensure robustness and cross-cohort reproducibility, the resulting gene sets from the discovery and validation cohorts were intersected, yielding conserved gene signatures for CAFs and pericytes.

### Bulk RNA-seq analysis and CAF quantification

The GSE190504 cohort [19] (n = 88: 23 oligodendrogliomas, 32 astrocytomas, 33 glioblastomas) was analyzed from the GEO-deposited processed expression matrix and accompanying series-matrix metadata (histology, IDH1 status, treatment). CAF activity was quantified in parallel by two complementary, transcriptome-scale read-outs: (i) a rank-based CAF signature score and (ii) a CAF cell-state proportion obtained by reference-based deconvolution with cellanneal [20]. All processing and testing were performed in Python (numpy, pandas, scipy.stats). The CAF signature (31 genes; 30 detected in GSE190504) was derived from our single-cell analysis (S2 Table). For each sample, all measured genes were ranked (average rank for ties) and rescaled to within-sample percentile ranks; the CAF rank score was then defined as the mean percentile rank of the CAF genes within that sample. This formulation captures the relative position of the CAF program within each transcriptome and is robust to library-size and cross-cohort distributional differences. A complementary CAF z-score (mean of per-gene standardized log2(expression + 1) across the CAF set) was computed as a sensitivity check. Group differences in the CAF rank and z scores across histology (oligodendroglioma, astrocytoma, glioblastoma) were assessed by an omnibus Kruskal–Wallis test followed by pairwise two-sided Mann–Whitney U (Wilcoxon rank-sum) tests, with Benjamini–Hochberg correction across the comparison family. For deconvolution, the 33 GBM bulk samples and an scRNA-seq–derived reference of 13 broad cell states (CAFs, the malignant subtypes AC-/MES1-/MES2-/NPC1-/NPC2-/OPC-like, GSCs, oligodendrocytes, endothelial cells, pericytes, T lymphocytes, TAMs) were harmonized to gene symbols (Ensembl version suffixes stripped, duplicated symbols collapsed by summation), intersected on the common gene space (∼14,950 genes after removal of rows zero in either matrix), and column-normalized to compositional vectors. cellanneal was run with disp_min = 0.5, bulk_min = 1e-8, bulk_max = 1, maxiter = 500; the bundled dual-annealing implementation was replaced by scipy.optimize.dual_annealing for NumPy ≥ 1.25 compatibility. Differences in the estimated CAF proportion between IDH1-wt and IDH1-mut, and between untreated and radiochemotherapy-treated GBM samples, were assessed by two-sided Mann–Whitney U tests. Concordance between the two CAF read-outs across the GBM subset was evaluated by Spearman rank correlation (Pearson reported as supportive). All tests were two-sided, with a BH-adjusted significance threshold of 0.05.

### Reference profiles for bulk deconvolution

Single-cell RNA-seq datasets from discovery and validation cohorts were integrated to generate reference expression profiles for bulk deconvolution. Cell-type annotations and patient metadata were harmonized across datasets, and low-quality cells and rare cell types (n < 50 cells) were excluded. Log-normalized gene expression values were used to compute average expression profiles for each annotated cell type. In addition, pseudo-bulk profiles were generated by aggregating cells within patient- and cell-type–specific groups to capture inter-sample variability. The resulting cell-type–level and pseudo-bulk reference matrices were used as input for downstream deconvolution analysis.

### Survival analysis

Survival analyses were performed using the online platform GEPIA3 [21], which integrates RNA-seq and clinical data from The Cancer Genome Atlas and Genotype-Tissue Expression Project. Analyses were conducted in the combined glioma cohort comprising lower-grade gliomas (LGG) and GBM. For evaluation of the CAF gene signature, all 31 genes included in the final cross-cohort CAF signature were entered simultaneously into the multigene survival module. For ligand–receptor interactions, survival analysis was performed using the complete set of genes constituting each interaction, including both ligand and receptor genes (and all receptor subunits when applicable). Patients were stratified into high- and low-expression groups according to the default GEPIA3 settings, and hazard ratios (HRs) and log-rank p-values were used to assess associations with overall and progression-free survival.

## Results

### CAFs constitute a distinct stromal population in glioblastoma and likely originate from pericytes

We first analyzed the discovery cohort, which integrated single-cell RNA-sequencing data from ten patients with primary IDH-wildtype GBM, including two recurrent tumors, together with patient-derived organoids (PDOs) and brain tumor-initiating cell (BTIC) lines. After quality control and integration, the dataset comprised 70,931 cells, including 255 cells annotated as cancer-associated fibroblasts (CAFs). CAFs were detected in all ten patient samples, indicating that this population is a consistent, albeit relatively rare, component of the glioblastoma microenvironment (Fig 1A, B; Fig S1). Differential gene expression, functional enrichment, and UCell analyses demonstrated that CAFs are characterized by a distinct transcriptional program enriched for genes involved in extracellular matrix organization, collagen fibril formation, and inflammatory responses (Fig 1C). Full term names, identifiers, and gene counts are provided in Fig S2. In contrast, pericytes and endothelial cells exhibited enrichment of their respective lineage-specific signatures, confirming the molecular distinction between these closely related vascular-associated populations. Together, these findings indicate that CAFs represent a transcriptionally unique stromal population with both structural and immunomodulatory properties within the GBM microenvironment. Diffusion map embedding and pseudotime analysis revealed a continuous transcriptional trajectory extending from ECs through pericytes to CAFs (Fig 1D, E; interactive 3D visualization available in the GitHub repository). This finding suggests that CAFs are transcriptionally linked to vascular-associated stromal populations and supports the hypothesis that they may arise through progressive phenotypic transition rather than representing an entirely independent lineage. Analysis of the transition probability matrix and generalized additive model (GAM) of gene expression dynamics along pseudotime provided additional support for this model (Fig 1F, G; Fig S3). The close transcriptional similarity between pericytes and CAFs, together with gradual and coordinated changes in gene expression along the trajectory, indicated a smooth transition between these two cell states rather than a sharp boundary.

**Fig 1.**
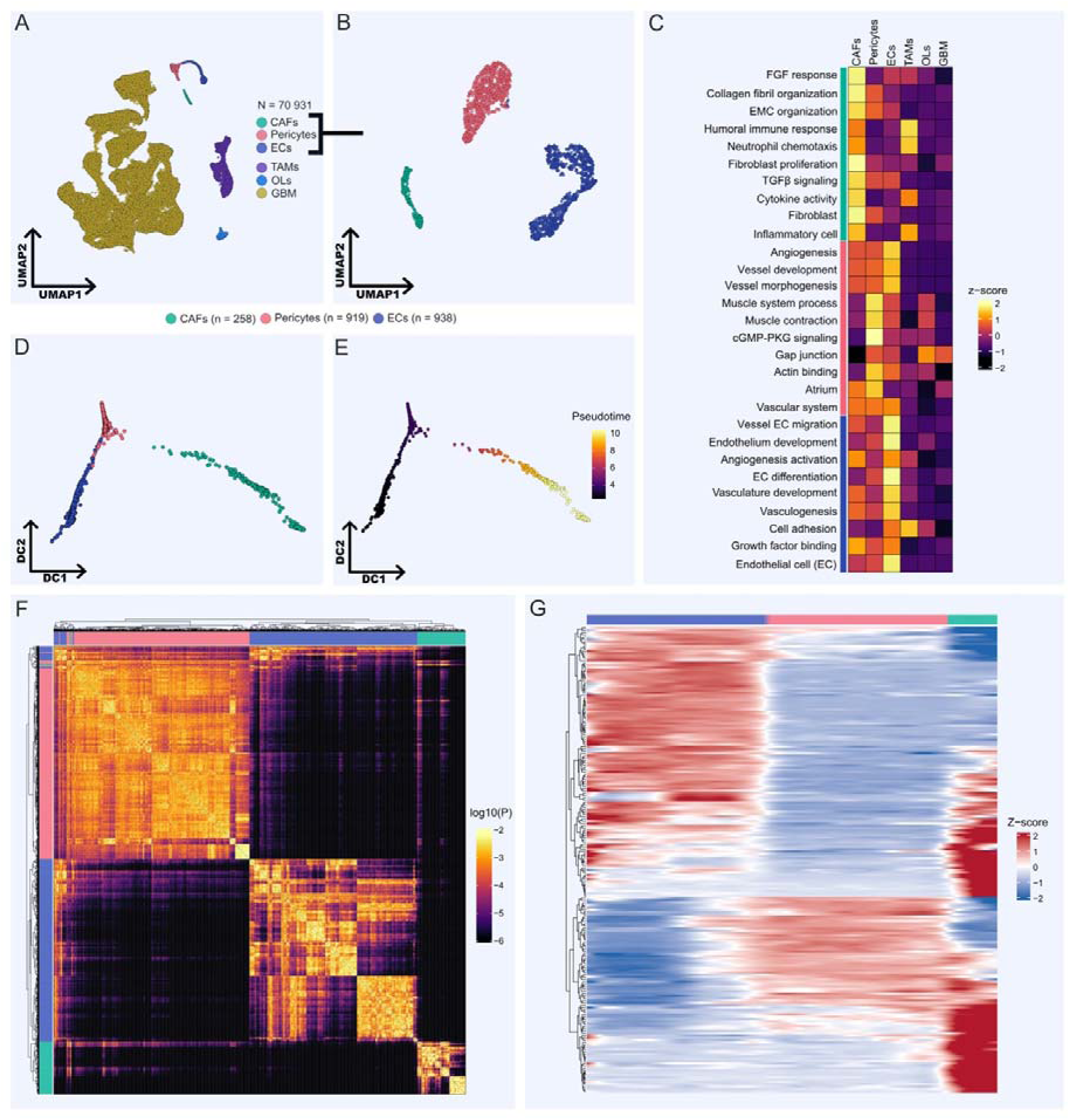
Discovery cohort. (A) Uniform manifold approximation and projection (UMAP) of the integrated discovery cohort comprising 70,931 cells from 10 patients with primary IDH-wildtype glioblastoma, including two recurrent tumors, together with patient-derived organoids and brain tumor-initiating cell lines. Major annotated cell populations include cancer-associated fibroblasts (CAFs, green), pericytes (red), endothelial cells (ECs, blue), tumor-associated macrophages/microglia (TAMs, purple), oligodendrocytes (OLs, bright blue), and glioblastoma cells (GBM, yellow). (B) UMAP representation restricted to CAFs, pericytes, and endothelial cells. (C) UCell enrichment analysis of Gene Ontology (GO) terms identified from the top 50 differentially expressed genes in CAFs, pericytes, and endothelial cells. For each cell type, significantly enriched GO terms were converted into corresponding gene sets and scored across all cells using UCell. Vertical bars indicate the cell type in whose differentially expressed gene set each GO term was significantly enriched (green, CAFs; red, pericytes; blue, endothelial cells). (D) Diffusion map embedding of CAFs, pericytes, and endothelial cells, demonstrating a continuous transcriptional continuum from endothelial cells through pericytes to CAFs. (E) Diffusion pseudotime projected onto the diffusion map embedding. Cells are colored according to pseudotime values, revealing a gradual progression that follows the same transcriptional continuum observed in panel D, from endothelial cells through pericytes to CAFs. (F) Transition probability matrix derived from diffusion map analysis. Three major transcriptional compartments corresponding to endothelial cells, pericytes, and CAFs are clearly distinguishable. Endothelial cells and CAFs display internal substructure, reflected by pronounced diagonal patterns and multiple subclusters, whereas pericytes form a more homogeneous compartment with comparatively limited internal heterogeneity. Although CAFs constitute a distinct cluster, transition probabilities indicate greater transcriptional similarity to pericytes than to endothelial cells. (G) Generalized additive model (GAM) of gene expression dynamics along pseudotime, illustrating gradual and coordinated transcriptional changes consistent with the transition from pericytes to CAFs.

To validate these findings, we analyzed an independent cohort comprising 44 glioblastoma samples generated across four separate scRNA-seq studies. After quality control and integration, the validation dataset contained 59,228 cells. CAFs were identified in samples from all four studies (Fig S4). In contrast to the discovery cohort, CAFs and pericytes formed a shared mesenchymal compartment on the global UMAP representation (Fig 2A). However, subclustering of this compartment readily resolved two transcriptionally distinct populations corresponding to pericytes and CAFs (Fig 2B). This observation further supports the close molecular relationship between these cell types while confirming that CAFs retain a rare and distinct transcriptional identity. Consistent with the discovery cohort, differential gene expression and functional enrichment analyses demonstrated that CAFs were enriched for genes associated with extracellular matrix organization, collagen fibril formation, and angiogenesis, whereas pericytes and endothelial cells displayed their expected lineage-specific transcriptional programs (Fig 2C, Fig S2). Diffusion map and pseudotime analyses recapitulated the same continuous transcriptional trajectory observed in the discovery cohort, extending from endothelial cells through pericytes to CAFs (Fig 2D, E; interactive 3D visualization available in the GitHub repository). Analysis of the transition probability matrix revealed extensive transcriptional connectivity among these populations, with the strongest non-self-transition probabilities observed between CAFs and pericytes (Fig 2F). Generalized additive model (GAM) analysis of gene expression dynamics along pseudotime similarly demonstrated a gradual shift from pericyte-associated to CAF-specific transcriptional programs (Fig 2G). Taken together, the validation cohort reproduced all major findings from the discovery analysis, providing strong independent support for the conclusion that CAFs represent a distinct stromal population in GBM and likely arise from pericytes.

**Fig 2.**
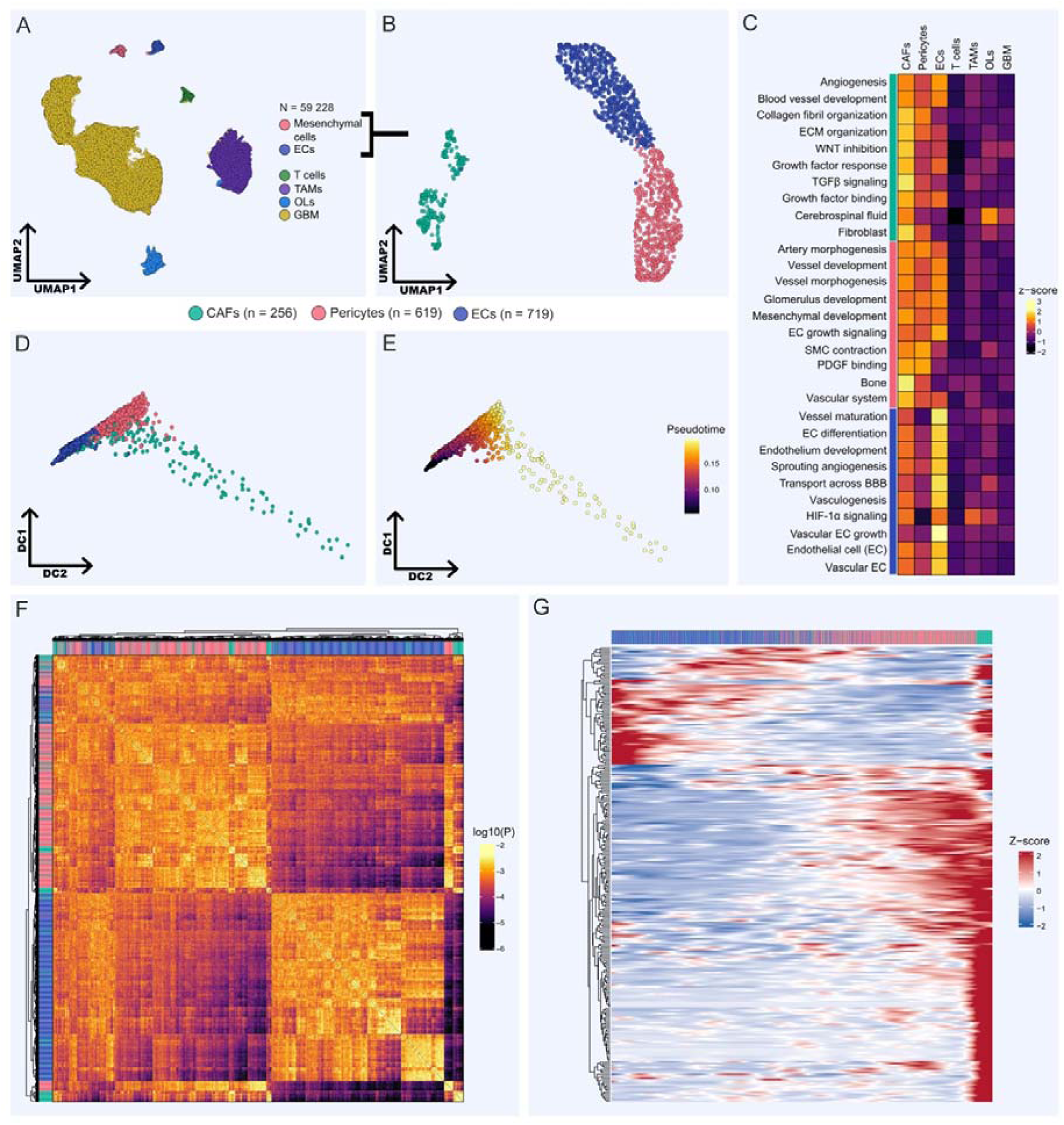
Validation cohort. (A) UMAP of the integrated validation cohort comprising 59,228 cells from 44 independent samples, including 36 IDH-wildtype glioblastomas and 8 high-grade gliomas. Major annotated cell populations include mesenchymal cells (comprising CAFs and pericytes; red), endothelial cells (ECs, blue), T lymphocytes (T cells, dark green), tumor-associated macrophages/microglia (TAMs, purple), oligodendrocytes (OLs, light blue), and malignant glioma cells (GBM, yellow). (B) UMAP representation restricted to CAFs, pericytes, and endothelial cells. Subsetting these populations resolves the mesenchymal compartment into two transcriptionally distinct clusters corresponding to CAFs and pericytes. (C) UCell enrichment analysis of Gene Ontology (GO) terms identified from the top 50 differentially expressed genes in CAFs, pericytes, and endothelial cells. For each cell type, significantly enriched GO terms were converted into corresponding gene sets and scored across all cells using UCell. Vertical bars indicate the cell type in whose differentially expressed gene set each GO term was significantly enriched (green, CAFs; red, pericytes; blue, endothelial cells). (D) Diffusion map embedding of CAFs, pericytes, and endothelial cells, demonstrating a continuous transcriptional continuum from endothelial cells through pericytes to CAFs, consistent with the pattern observed in the discovery cohort. (E) Diffusion pseudotime projected onto the diffusion map embedding. Cells are colored according to pseudotime values, revealing a gradual progression that follows the same transcriptional continuum observed in panel D. (F) Transition probability matrix derived from diffusion map analysis. A large mixed compartment composed predominantly of CAFs and pericytes occupies the upper-left portion of the matrix and is characterized by high transition probabilities between these two populations. Endothelial cells form a largely distinct compartment with limited overlap with mesenchymal cells. Two smaller clusters corresponding to more differentiated pericytes and CAFs remain connected by elevated transition probabilities. Consistent with the discovery cohort, CAFs exhibit greater transcriptional similarity to pericytes than to endothelial cells. (G) Generalized additive model (GAM) of gene expression dynamics along pseudotime, illustrating gradual and coordinated transcriptional changes consistent with the transition from pericytes to CAFs, recapitulating the pattern observed in the discovery cohort.

### CAF’s cell-cell communication network

To identify robust CAF-associated intercellular interactions, we focused on ligand– receptor pairs that were significantly detected in both the discovery and validation cohorts. For cell–cell communication analysis, the malignant GBM compartment was further subclustered into established transcriptional states, including MES1-like, MES2-like, NPC1-like, NPC2-like, OPC-like, AC-like, and GSCs [1, 15, 22]. This approach yielded 17 reproducible interactions, representing a conserved core communication network involving CAFs within the GBM microenvironment (Fig 3A, B). To prioritize interactions with potential clinical relevance, we evaluated the prognostic impact of each ligand–receptor pair by calculating hazard ratios (HRs) based on Kaplan–Meier survival analysis using the combined expression of all genes constituting each pair (Fig 3C). Several interactions were strongly associated with poor survival (HR > 4), including signaling through integrin receptors (NAMPT→α5β1, ANGPTL4→α5β1, and FBN1→α5β1), as well as SERPINE1→PLAUR, MIF→CD44+CD74, COL1A1→CD44 and ADM→CALCRL mediated pathways. CAFs emerged as a major source of several high-risk ligands, including COL1A1, FBN1, and SERPINE1 (Fig 3D, E). These molecules are implicated in key tumor-promoting processes. Signaling mediated by integrin receptors and ADM→CALCRL can support endothelial activation and angiogenesis [23–26]. Interactions involving CD44 and CD74 have been linked to immune suppression [27–30]. Furthermore, COL1A1→CD44 signaling may contribute to the maintenance of stemness-associated properties in cancer cells, consistent with the established role of CD44 as a surface marker of cancer stem cells [31, 32]. Among the reproducible interactions, SERPINE1→LRP2 was notable for its specificity to CAF→oligodendrocyte communication in both cohorts. Although this interaction was associated with an elevated hazard ratio, its prognostic effect appeared to be driven primarily by *SERPINE1* expression, as *LRP2* expression alone did not show association with survival (HR = 1). The full list of ligand–receptor interactions identified in both the discovery and validation datasets is provided in Table S1. Overall, the conserved ligand–receptor interactions identified across independent cohorts suggest that CAFs participate in signaling networks involved in angiogenesis, extracellular matrix remodeling, and immune modulation.

**Fig 3.**
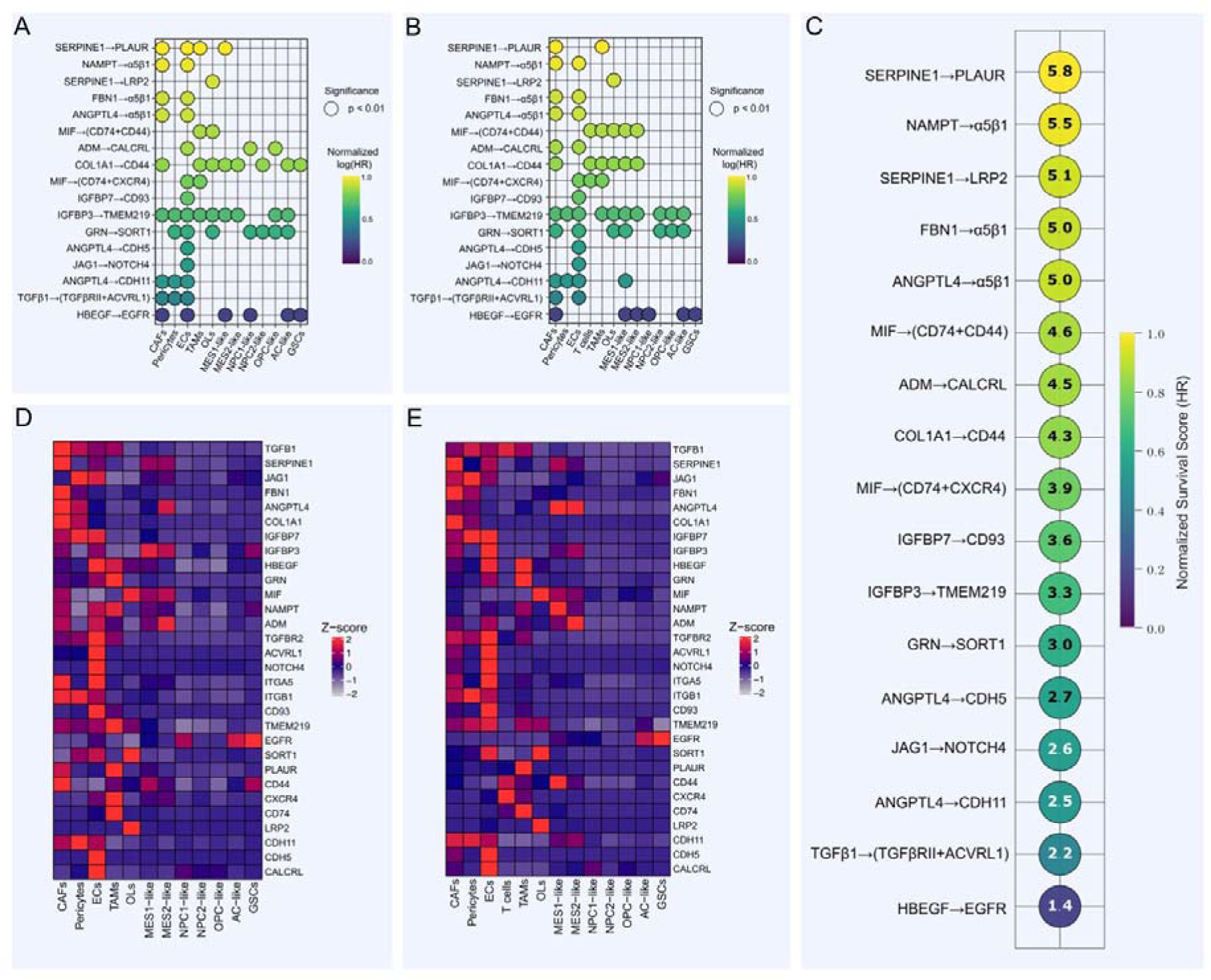
CAF-associated cell–cell communication landscape. (A-B) Outgoing signaling pathways from CAFs in the discovery (A) and validation (B) cohorts. Only ligand–receptor interactions reproduced in both datasets are shown. Pathways are colored according to normalized hazard ratio (HR) values derived from Kaplan–Meier survival analysis based on the co-expression of genes comprising each ligand–receptor pair. Interactions are ordered by their association with patient survival, from the strongest to the weakest adverse effect. (C) Hazard ratio estimates for each reproducible ligand–receptor interaction shown as absolute HR values. Interactions are ranked according to their impact on overall survival, from the highest to the lowest HR. For visualization purposes, HR values were normalized to the logarithm of the maximum observed HR value (HR = 5.8). (D-E) Heatmaps of expression for all genes involved in the selected ligand–receptor interactions in the discovery (D) and validation (E) cohorts. Overall, both heatmaps show highly consistent expression patterns across the two independent datasets.

### CAF transcriptional signature reveals association with poor prognosis and therapy-related expansion

To derive a robust CAF-specific gene signature, we identified genes that were consistently enriched in CAFs in both the discovery and validation cohorts (see Methods). This analysis yielded a 31-gene signature comprising canonical fibroblast markers and genes associated with activated stromal states, including *FAP*, *COL1A1*, *COL6A2*, *COL6A3*, *FBN1*, *LTBP1*, and *FBLN2* (Fig 4A, B). Functional enrichment analysis confirmed strong association with extracellular matrix organization, TGF-β signaling, growth factor binding, and fibroblast-related transcriptional programs (Table S2). Enrichment of the “liver epithelial cells” term was primarily driven by *CYP1B1* and *NBL1*, with *CYP1B1* showing consistent overexpression in CAFs across both cohorts. Co-expression of the CAF signature was strongly associated with adverse clinical outcomes in the combined lower-grade glioma and glioblastoma cohort (LGG-GBM), with significant associations observed for both overall survival (hazard ratio [HR] = 4.62) and progression-free survival (HR = 2.99) (Fig 4C). These findings indicate that transcriptional programs associated with CAFs are linked to aggressive disease behavior. To further investigate the clinical relevance of CAFs, we analyzed a bulk RNA-seq cohort comprising 88 glioma samples (including glioblastoma, astrocytoma, and oligodendroglioma) with available data on histology, *IDH1* mutation status, and treatment (RCT – radio-chemotherapy). CAF abundance was estimated using two complementary approaches: rank-based scoring of the CAF signature across all samples and bulk deconvolution in glioblastoma samples with sufficient representation of stromal components (n = 33). CAF signature scores differed significantly across histological subtypes, with the highest values observed in glioblastoma compared with astrocytoma and oligodendroglioma (Kruskal–Walli’s test, BH-adjusted p = 0.001) (Fig 4D). Both rank-based scoring and deconvolution consistently demonstrated increased CAF expression and abundance in IDH-wildtype tumors relative to IDH-mutant gliomas (Fig 4D–F), with statistically significant differences observed by both methods. Analysis of treatment status revealed a similar pattern. In the full glioma cohort, rank-based CAF scores did not differ significantly between treated and untreated tumors. However, when analysis was restricted to glioblastoma samples, treated tumors exhibited significantly higher CAF scores, consistent with the results of bulk deconvolution, which also showed a significant increase in CAF proportions following therapy (Fig 4E, F). The two approaches showed strong concordance, with a Spearman correlation coefficient of 0.82 (p = 4.54 × 10 ^9^’; Fig 4G). The CAF and pericyte signature gene lists are provided in Table S3, while the full table of statistical test results is available in Table S4.

**Fig 4.**
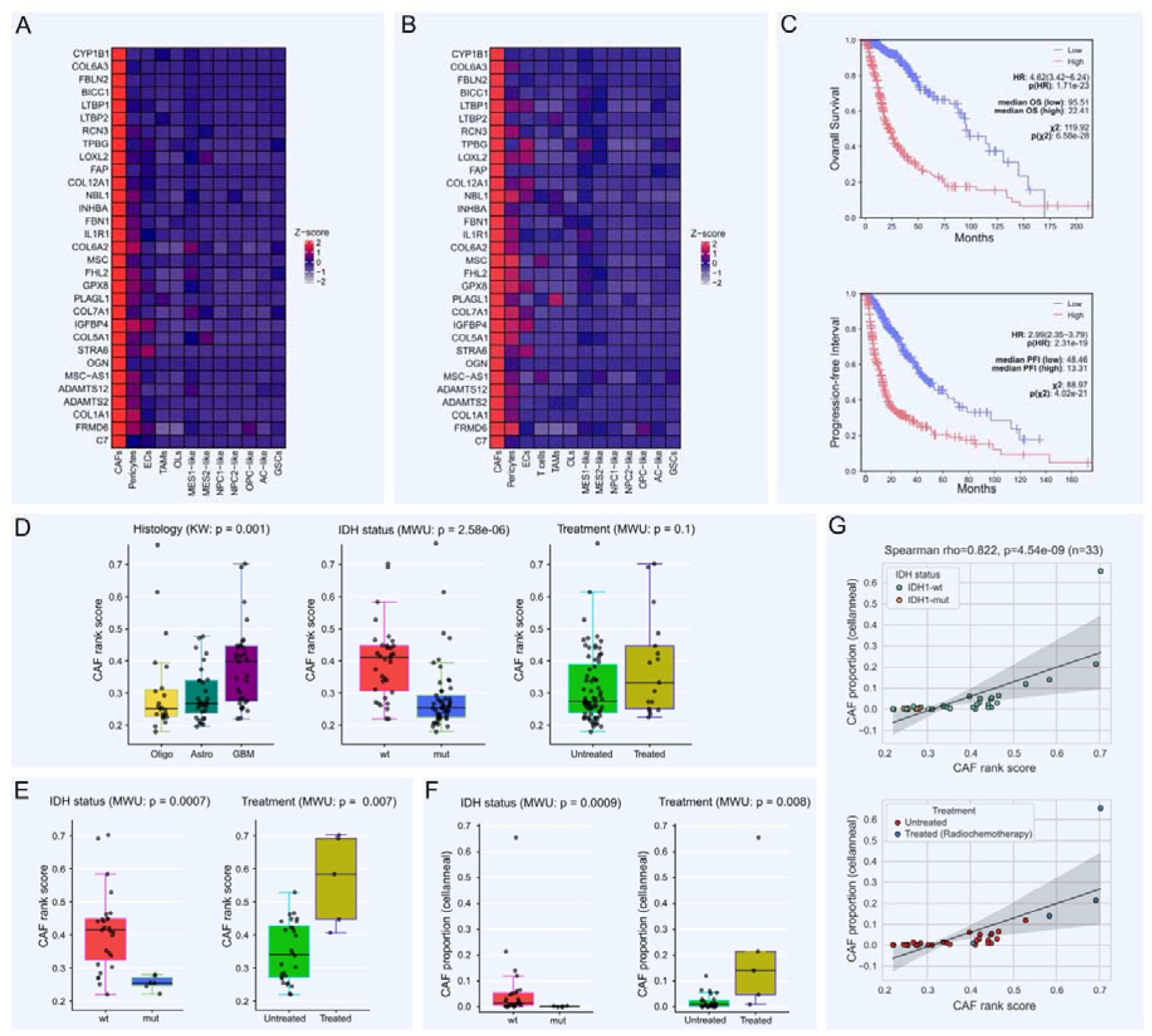
CAF signature and bulk RNA-seq analysis of CAF dynamics across glioma states. (A-B) Heatmaps showing the expression of CAF-specific genes identified by iterative differential expression analysis in the discovery and validation single-cell RNA-seq cohorts. (C) Kaplan–Meier analysis of the CAF signature in a combined cohort of lower-grade glioma and glioblastoma patients (LGG + GBM) showing that high CAF signature expression is associated with shorter overall survival (OS; HR = 4.62) and progression-free survival (PFS; HR = 2.99). (D) Rank-based expression analysis of the CAF signature in the full glioma cohort (left; LGG + GBM; n = 88). CAF signature expression differed significantly among glioblastoma (WHO grade IV), astrocytoma (WHO grade II), and oligodendroglioma (WHO grade II) samples (Kruskal–Wallis test, BH-adjusted p = 0.0013). Expression was significantly higher in IDH-wildtype compared with IDH-mutant gliomas (Mann–Whitney U test, p = 1.03 × 10□□). No significant difference was observed between treatment-naïve and post-radiochemotherapy (RCT) samples in the full cohort (right panel, Mann–Whitney U test, p = 0.10). (E) Rank-based CAF signature expression analysis restricted to glioblastoma samples (n = 33). CAF signature expression remained significantly higher in IDH-wildtype compared with IDH-mutant tumors (left; Mann–Whitney p = 0.0007) and was increased in post-radiochemotherapy samples relative to treatment-naïve tumors (right; Mann–Whitney p = 0.004). (F) Bulk deconvolution analysis using Cellanneal in the glioblastoma cohort (n = 33). Estimated CAF abundance was significantly higher in IDH-wildtype tumors than in IDH-mutant tumors (left; Mann–Whitney p = 0.0009) and in post-radiochemotherapy samples compared with treatment-naïve tumors (right; Mann–Whitney p = 0.008). (G) Concordance between rank-based CAF signature scores and deconvolution-derived CAF abundance estimates in glioblastoma samples, demonstrating a strong positive correlation (Spearman’s ρ = 0.82, p = 4.54 × 10□^9^).

Collectively, these results demonstrate that a reproducible CAF transcriptional signature is associated with poor prognosis and reveal that CAF abundance is increased in IDH-wildtype and treatment-exposed glioblastomas, supporting a role for CAFs in therapy-associated remodeling of the TME.

## Discussion

In various types of malignant tumors, cancer-associated fibroblasts (CAFs) are a well-characterized stromal component and key regulators of tumor progression [33]. However, in the context of GBM, their existence and functional relevance have long remained a subject of skepticism. Only in recent years has interest in CAF-like populations in gliomas increased [3, 4, 6, 34, 35], most investigations have primarily addressed their immunosuppressive functions and direct pro-tumorigenic effects on glioblastoma cells. At the same time, a robust and reproducible CAF signature for GBM remains unavailable, largely due to the substantial transcriptional similarity between CAFs and pericytes, which complicates their clear separation and may confound the identification of a genuine CAF-specific transcriptional profile.

One possible reason for this limited attention to CAFs in neuro-oncology is the technical difficulty of detecting these cells in single-cell RNA-seq datasets, as tumor-associated cell populations (including CAFs) may be poorly preserved during library preparation. Similar limitations also apply to other components of the GBM microenvironment, including pericytes, endothelial cells, astrocytes, and neurons. A key factor is the enzymatic tissue dissociation step, which can lead to damage or loss of sensitive cell populations [7]. As a result, certain cell types may undergo selective depletion and fail to be captured in single-cell data, particularly in the case of rare populations. Additional explanations include the potentially low baseline abundance of CAFs in the glioma stroma and their highly context-dependent nature, suggesting that their emergence or detection may be tightly linked to specific pathophysiological conditions rather than being a constant feature across all samples. These factors are not mutually exclusive and likely act in concert. Nevertheless, even the sporadic detection of CAFs in GBM holds critical biological significance, as it may reflect specific cellular transformations occurring within the tumor and serve as an indicator of pronounced changes in the TME that give rise to these cells. At the same time, the precise triggers driving these phenotypic shifts and the subsequent functional role of CAFs in gliomas remain open questions.

The cellular origin of cancer-associated fibroblasts (CAFs) remains a subject of ongoing debate, with various potential sources described in the literature, including quiescent fibroblasts, endothelial cells, pericytes, stellate cells, bone marrow-derived mesenchymal cells, adipocytes, and glial cells [1, 8–13]. Within the specific context of GBM, we hypothesize that local pericytes and endothelial cells represent the two most plausible candidate progenitor populations due to their abundance and plasticity within the TME. Alternative origins have been proposed, but they often imply less parsimonious mechanisms. For instance, Lootens et al. suggested that CAFs arise from tumor-associated mesenchymal stem/stromal cells (TA-MSCs), presenting a multi-step process that requires the recruitment of bone marrow-derived MSCs via the vasculature, their homing, and subsequent sequential differentiation into TA-MSCs and then CAFs [34]. From our perspective, such a systemic recruitment mechanism appears unnecessarily complex compared to the localized activation of resident vascular cells. In another study, the endothelial-to-mesenchymal transition (EndMT) was highlighted as the primary driver of CAF generation in glioblastoma [36]. While EndMT remains a viable pathway, it involves a fundamental transdifferentiation across distinct lineages. In contrast, the pericyte-to-fibroblast transition (PFT), primarily driven by localized PDGF-BB/PDGFRβ signaling, offers a more direct and biologically efficient route [12, 37]. This vascular-centric model is strongly supported by spatial tissue architecture; for instance, Saket Jain et al. explicitly identified CAFs within the perivascular niche, in close proximity to pericytes and endothelial cells, while maintaining a distinct cellular identity [6]. Crucially, our own analysis provides computational support for this spatial alignment. By evaluating two independent single-cell RNA-seq cohorts, we identified a robust and reproducible transcriptional trajectory directly capturing the transition from pericytes to CAFs. Given these convergent findings, combined with their shared mesenchymal lineage and baseline transcriptional similarity, we propose that resident pericytes serve as the primary progenitor population driving CAF heterogeneity in GBM.

Based on the cell–cell communication analysis, it is difficult to draw definitive conclusions regarding the functional role of CAFs in glioblastoma. Nevertheless, we identified a set of reproducible ligand–receptor interactions associated with angiogenesis and immune modulation that may reflect biologically relevant signaling pathways. Although the involvement of CAFs in immunosuppression and angiogenesis has been reported previously in glioblastoma studies [6, 34, 38], our analysis identified reproducible CAF-associated immunosuppressive and pro-angiogenic interactions with particularly strong prognostic impact (HR > 4), many of which have not previously been described in the context of GBM. The SERPINE1→PLAUR interaction merits particular attention because of its strong association with adverse survival outcomes. Although its specific biological role in GBM remains unclear, PLAUR/uPAR expression has previously been identified in glioblastoma-associated macrophages, where it has been linked to a protumorigenic TME and proposed as a potential therapeutic target [39]. Similarly, while MIF activity has been linked to immunosuppressive remodeling of the tumor microenvironment [40, 41], CAF-derived MIF-mediated immunosuppression has not previously been described in glioblastoma. Regarding the potential pro-angiogenic effects of CAFs identified in this study, CAF-derived ligands such as NAMPT, ANGPTL4, FBN1, and ADM have also not been previously reported in glioblastoma. Given the pleiotropic nature of the corresponding receptors, their ability to interact with ligands produced by multiple cell types, and the relatively low abundance of CAFs in our datasets, the extent to which CAFs quantitatively contribute to angiogenesis and immune suppression in GBM remains uncertain. Nonetheless, the reproducibility of these interactions across two independent cohorts suggests that CAFs participate in signaling networks associated with key tumor-promoting processes.

Interestingly, pericyte-derived CAFs were significantly enriched in IDH1-wildtype gliomas. Given that IDH1-wildtype status together with TERT promoter mutations represent the molecular hallmarks of glioblastoma and are consistently associated with poor clinical outcomes [42, 43], this association suggests that CAFs are preferentially enriched within the most aggressive glioma subtype. This observation is consistent with the pro-tumorigenic functions of CAFs identified in our study and further supports their potential contribution to the aggressive biology of glioblastoma.

The contribution of CAFs to chemoresistance is well-documented across various solid tumors [44–47]; however, in the specific context of GBM, literature linking tumor-associated fibroblasts to temozolomide (TMZ) resistance remains sparse. To date, only a limited number of studies have addressed this axis: Zuo et al. attributed resistance to CAF-secreted CCL2 cytokine [35], while Li Ji et al. implicated TNC and FLNC as drivers of both EndMT-mediated CAF generation and subsequent TMZ resistance [36]. This gap in understanding is particularly critical given the classic clinical paradox of GBM: the therapeutic benefit of standard-of-care chemoradiotherapy is almost universally confined to the initial phase of treatment, after which the tumor rapidly develops drug resistance, leading to inevitable progression. This transition is driven by therapy-induced remodeling of the GBM microenvironment, though the exact underlying mechanisms have yet to be fully elucidated. Our findings offer a novel perspective on this phenomenon. We first observed a progressive increase in CAF abundance and a marked enrichment of the CAF transcriptional signature in post-treatment GBM samples. Crucially, CAFs exhibited highly selective expression of *CYP1B1* a member of the cytochrome P450 superfamily historically associated with resistance to multiple chemotherapeutic agents in breast, renal, and other cancers [48–51], but whose direct link to temozolomide metabolism in GBM has not been previously reported.

Taken together, these observations allow us to hypothesize that CAFs in glioblastoma may actively emerge during therapy-induced remodeling of the TME. Because our analysis indicates a pericyte-derived origin for these cells, they are inherently positioned within the perivascular space, directly adjacent to the capillary bed. This strategic localization at the blood-tumor interface, combined with their selective expression of *CYP1B1*, suggests a potential functional role: these pericyte-derived CAFs might act as a localized metabolic barrier, participating in the detoxification or clearance of temozolomide before it reaches the deeper tumor parenchyma. Consequently, although CAFs represent a relatively minor cell population in GBM compared to other solid tumors, their enrichment after treatment point toward a specialized pathophysiological response aimed at maintaining chemoresistance. While these observational findings are inherently predictive and require rigorous experimental validation to define the exact kinetics of CYP1B1-mediated drug metabolism, they highlight pericyte-to-CAF transition as a plausible mechanism contributing to the post-treatment evolution of the GBM microenvironment.

## Acknowledgements

The authors would like to sincerely thank Lidia Garkul for her continuous support throughout the preparation of this manuscript, for valuable discussions, insightful suggestions, and for accompanying the development of this project from its early stages. We are also grateful to Anastasia Mikhailova and Artem Burtsev for their help in introducing and guiding us through single-cell RNA sequencing analysis and related concepts. Finally, we thank Dr. Nisan for the valuable suggestion to incorporate bulk RNA-seq deconvolution analysis, which significantly strengthened the study. We thank Aldo Spallone for insightful discussions and valuable scientific ideas at the early stages of this work.

## Author Contributions

AI: developed the study concept, performed data analysis and code development, and contributed to manuscript writing.

MP: principal investigator; supervised the project, contributed to study design, and participated in manuscript writing and critical revision.

## Data and Code Availability

The discovery cohort was based on the Gene Expression Omnibus (GEO) dataset GSE173278. The validation cohort was generated by integrating the publicly available GEO datasets GSM3828672, GSE103224, and GSE135045, together with the publicly available 10x Genomics GBM 3′ v3 scRNA-seq dataset. Bulk RNA-seq data were obtained from the GEO dataset GSE190504. All code used for data processing, analysis, figure generation, and 3D diffusion map visualization is publicly available at: https://github.com/neuropromotion/CAFs-in-glioblastoma-microenvironment.

## Supporting information

**Fig S1. Overview of the discovery single-cell RNA sequencing cohort.** (A) UMAP visualization of the full discovery dataset colored by annotated cell clusters. (B) Subannotation of glioblastoma cells into transcriptional subtypes. (C) Distribution of CAFs, pericytes, and endothelial cells across the 10 samples included in the discovery cohort. (D) Heatmap showing the top five differentially expressed genes for each annotated cluster. (E) Distribution of cells across samples in the discovery dataset.

**Fig S2. Functional enrichment and UCell analysis of CAF-, pericyte-, and endothelial cell-associated transcriptional programs.** (A) Discovery cohort. The top 50 differentially expressed genes for CAFs, pericytes, and endothelial cells (ECs) were subjected to functional enrichment analysis. Gene sets from significantly enriched terms (BH-adjusted p < 0.05) were subsequently used for UCell scoring across all annotated clusters. Heatmaps show cluster-level enrichment patterns for the resulting functional programs. (B) Validation cohort. UCell analysis was performed using the same workflow and gene sets derived from significantly enriched functional terms. Heatmaps demonstrate reproducible enrichment patterns across annotated clusters in the validation dataset.

**Fig S3. Dynamic gene expression changes along the endothelial cell–pericyte–CAF pseudotime trajectory.** (A) Discovery cohort. Heatmap showing generalized additive model (GAM)-based smooth expression variation across pseudotime along the endothelial cell (EC)– pericyte–CAF trajectory. Rows represent dynamically regulated genes and columns represent pseudotime progression. (B) Validation cohort. Heatmap showing GAM-based smooth expression variation across pseudotime along the EC–pericyte–CAF trajectory in the validation dataset. Gene names are indicated alongside the heatmaps.

**Fig S4. Overview of the validation single-cell RNA sequencing cohort.** (A) UMAP visualization of the full validation dataset colored by annotated cell clusters. (B) Subannotation of glioblastoma cells into transcriptional subtypes. (C) Distribution of CAFs, pericytes, and endothelial cells across samples included in the validation cohort. (D) Heatmap showing the top five differentially expressed genes for each annotated cluster. (E) Distribution of cells across samples in the validation dataset.

**Table S1. Reproducible ligand–receptor interactions identified by CellChat analysis within discovery and validation datasets.** This table contains statistically significant ligand–receptor interactions identified using the CellChat framework and retained for downstream analysis. Each row corresponds to one directed interaction between a sender (source) and receiver (target) cell population.

**Table S2. Functional enrichment analysis of the CAF gene signature.**

**Table S3. Specific signatures for CAFs and Pericytes.**

**Table S4. Bulk RNA-seq analysis and cancer-associated fibroblast (CAF) quantification in diffuse glioma using rank-based signature scoring and CellAnn deconvolution.** This supplementary table summarizes the quantification of cancer-associated fibroblasts (CAFs) across 88 bulk RNA-seq samples from the GSE190504 diffuse glioma cohort. CAF abundance was estimated using two complementary approaches: (i) a rank-based CAF signature score (CAF_rank_score) derived from the relative expression of CAF marker genes, and (ii) a deconvolution-based estimate (CAF_cellanneal_deconvolution) obtained using cellanneal with reference profiles derived from our single-cell RNA-seq analysis. The table includes six worksheets described below.

