## Supplementary figures and images for "Pericyte-Derived Cancer-Associated Fibroblasts Correlate with Poor Survival and Are Enriched After Chemoradiotherapy in Glioblastoma"

### Figure S1

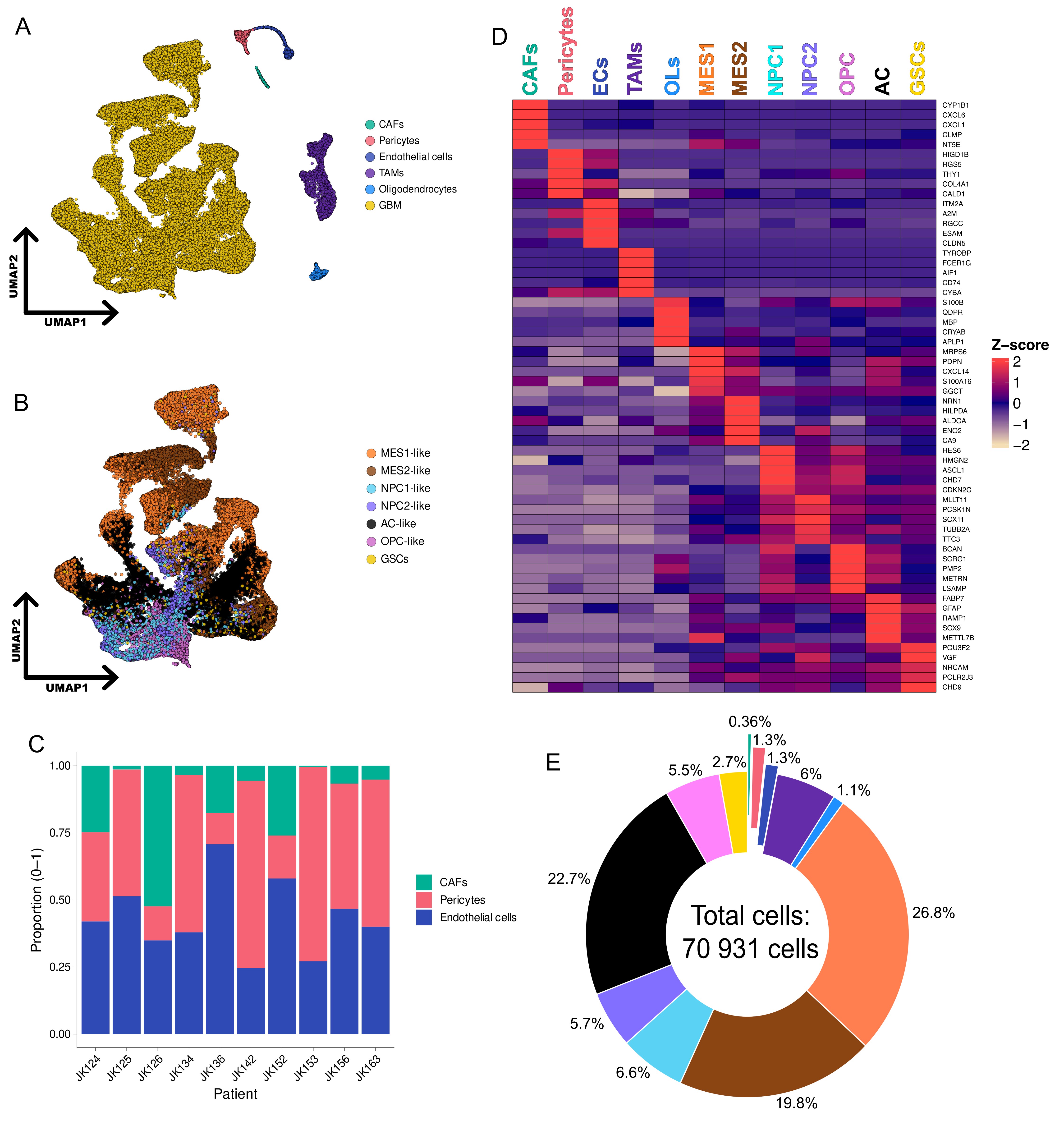

### Figure S2

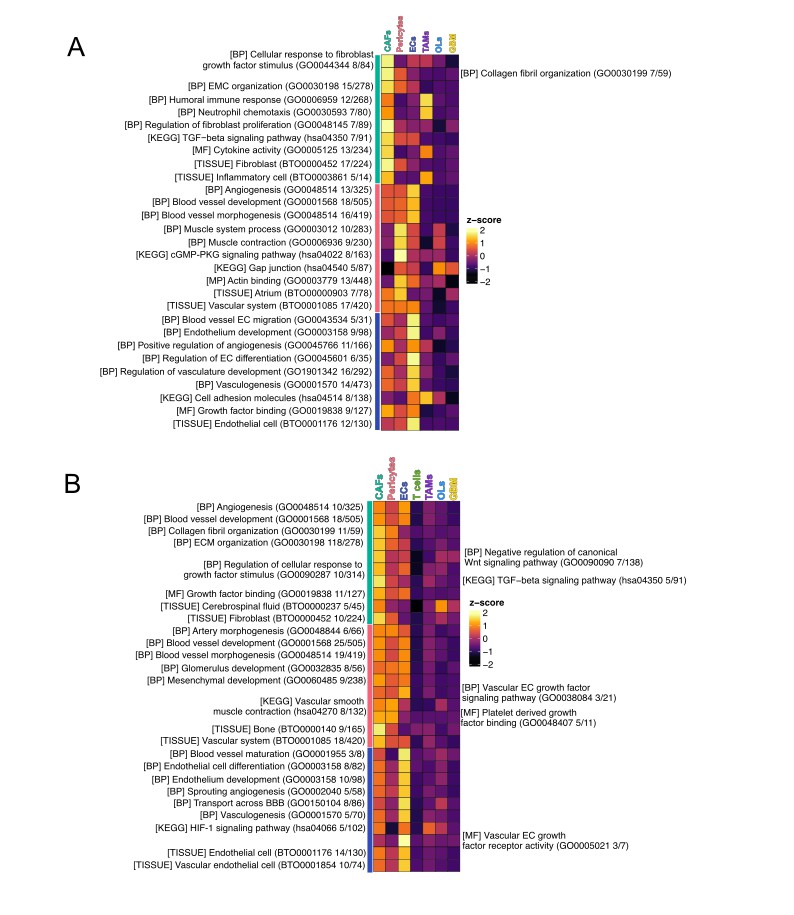

### Figure S3

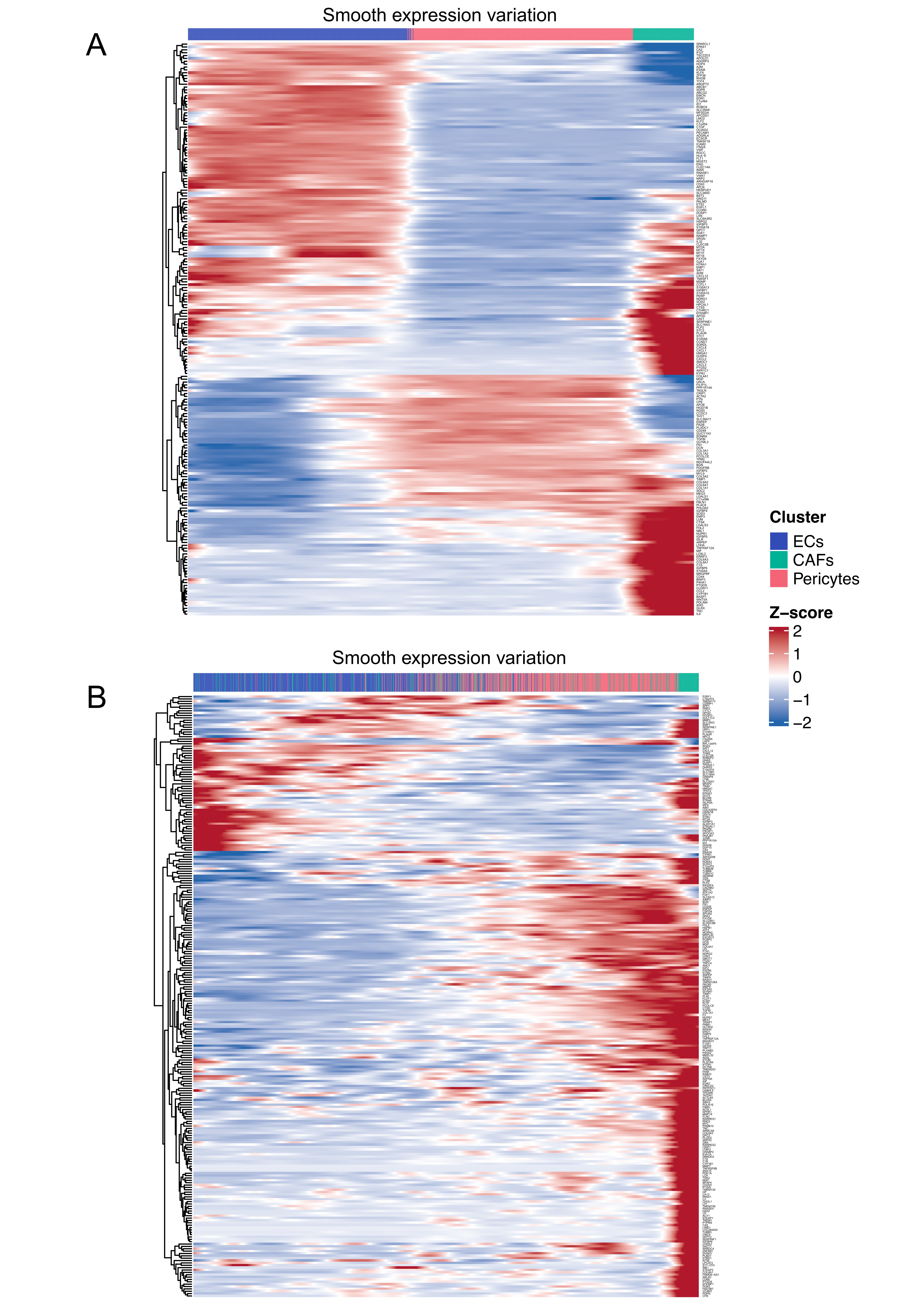

### Figure S4

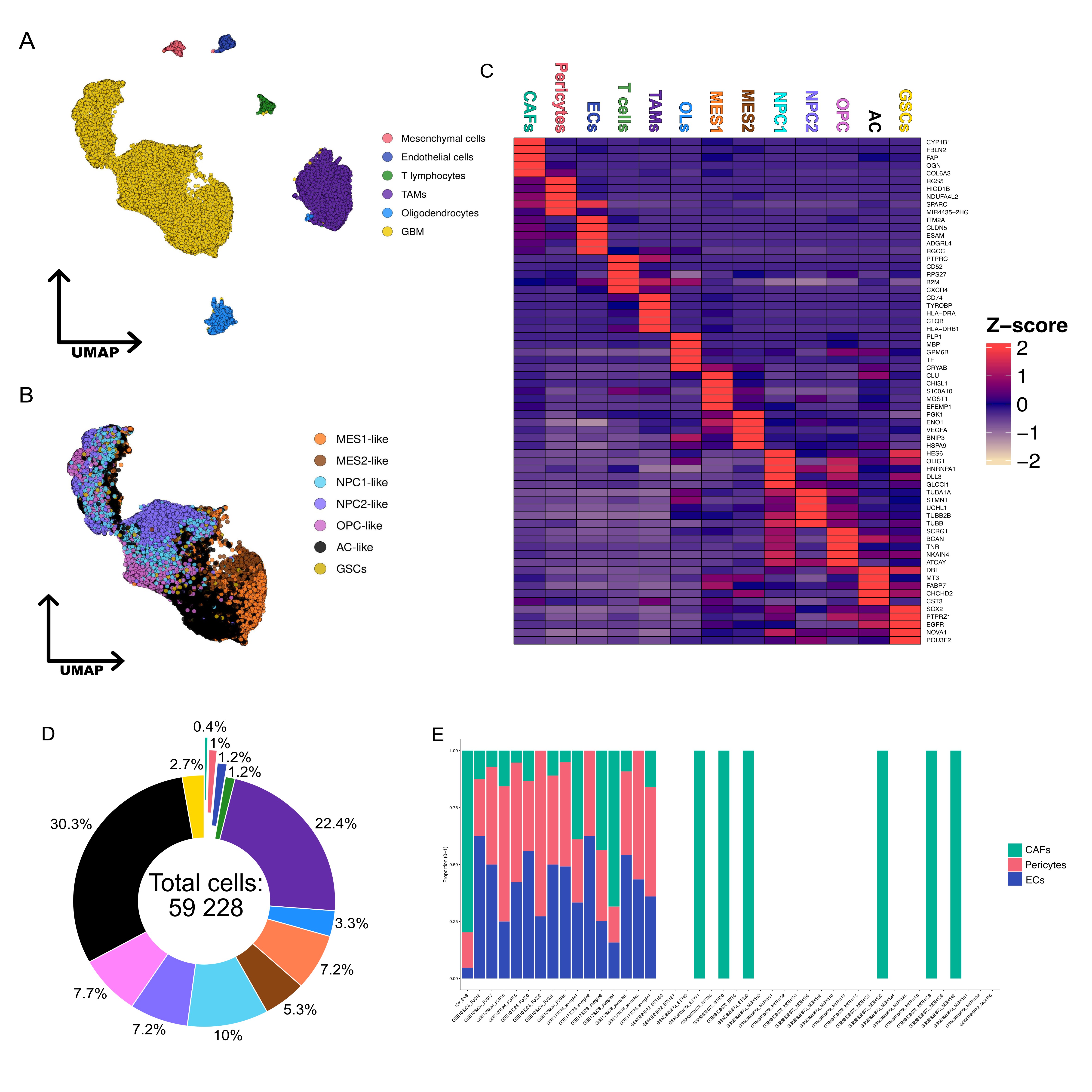
